# An adult brain atlas reveals broad neuroanatomical changes in independently evolved populations of Mexican cavefish

**DOI:** 10.1101/648188

**Authors:** Cody Loomis, Robert Peuß, James Jaggard, Yongfu Wang, Sean McKinney, Stephen Raftopoulos, Austin Raftopoulos, Daniel Whu, Matthew Green, Suzanne E. McGaugh, Nicolas Rohner, Alex C. Keene, Erik R. Duboue

## Abstract

A shift in environmental conditions impacts the evolution of complex developmental and behavioral traits. The Mexican cavefish, *Astyanax mexicanus*, is a powerful model for examining the evolution of development, physiology, and behavior because multiple cavefish populations can be compared to an extant and ancestral-like surface population of the same species. Many behaviors have diverged in cave populations of *A. mexicanus*, and previous studies have shown that cavefish have a loss of sleep, reduced stress, an absence of social behaviors, and hyperphagia. Despite these findings, surprisingly little is known about the changes in neuroanatomy that underlie these behavioral phenotypes. Here, we use serial sectioning to generate a brain atlas of surface fish and three independent cavefish populations. Volumetric reconstruction of serial-sectioned brains confirms convergent evolution on reduced optic tectum volume in all cavefish populations tested. In addition, we quantified volumes of specific neuroanatomical loci within several brain regions, which have previously been implicated in behavioral regulation, including the hypothalamus, thalamus, and habenula. These analyses reveal an expansion of the hypothalamus across all three cavefish populations relative to surface fish, as well as subnuclei-specific differences within the thalamus and habenulae. Taken together, these analyses support the notion that changes in environmental conditions are accompanied by neuroanatomical changes in brain structures associated with behavior. This atlas provides a resource for comparative neuroanatomy of additional brain regions and the opportunity to associate brain anatomy with evolved changes in behavior.

## Introduction

Shifts in environmental conditions drive evolutionary changes in development, morphology, and behavior (1–3). While the genetic basis of many behaviors has been studied extensively, much less is known about how changes in brain anatomy underlie behavioral evolution. Interspecies comparative approaches are often used to associate anatomical or neural circuit changes with evolved behavioral differences (4–6). However, these studies often focus on particular brain regions of interest and interpretations may be limited by the indirect nature of comparing different species. The generation of detailed anatomical brain atlases of individuals of the same species with divergent behavioral traits would provide insight into evolved changes in brain morphology that may associate with behavioral evolution.

The Mexican cavefish, *Astyanax mexicanus* provides the unique opportunity to investigate the relationship between brain anatomy and behavioral evolution in a single species (7–11). These fish exist as an eyed and pigmented population that inhabits the rivers of northeast Mexico and southern Texas, and at least 29 independent populations of largely blind and depigmented fish that inhabit the caves of northeast Mexico’s Sierra de El Abra and Sierra de Guatemala regions (Mitchell et al., 1977). Both surface and cave populations are interfertile, which allows for a direct comparisons of populations from the same species with different and well-described habitats and evolutionary history (13,14). Comparisons between surface fish and cavefish populations reveal evolved differences in diverse behavioral traits ranging from social behavior to sleep, and the emergence of these behaviors in multiple cavefish populations has established *A. mexicanus* as a model for convergent evolution (11,15–18).

A number of anatomical differences have been identified between surface fish and cavefish, including a reduction in brain regions associated with visual processing in cavefish and an expansion of the hypothalamus which is associated with social behavior (7,19,20). Nevertheless, *A. mexicanus* lacks a detailed brain atlas and little is known about the extent of anatomical changes between individual populations of cavefish. Further, an anatomical comparison between adult cave populations has not been performed, and it remains unclear if distinct or shared changes in brain anatomy underlie the behavioral differences observed between independently evolved cavefish populations.

Here, we have used serial sectioning of Nissl-stained brains, followed by volumetric reconstruction to generate a brain atlas for surface fish and three different populations of cavefish. Our analysis focuses on hypothalamic, thalamic, and habenular regions, which have previously been associated with behaviors known to diverge between surface fish and cavefish including responses to stress, social behavior, sleep regulation, feeding, and sensory processing (8,11,15–18). Our findings reveal an expansion of thalamic and habenular regions in cavefish, accompanied by a reduction in regions associated with visual processing. Strikingly, some subnuclei within the hypothalamus are expanded in cavefish, while other hypothalamic regions remain unchanged. Together, these findings provided a detailed anatomical reference for *A. mexicanus and* provide insight into the anatomical plasticity that accompanies the evolution of multiple behaviors.

## Results

### Volumetric reconstruction of serial sectioned adult brains

To generate an adult brain atlas, we serially sectioned brains of adult *A. mexicanus* from surface fish and three independent populations of cavefish: Pachón, Molino and Tinaja (Figure 1A). The Pachón and Tinaja populations are ‘old lineage’ and are closely related, while fish from the Molino population represent a ‘new lineage.’ All cave populations are thought to be independently derived origins of the cave phenotype (21–23). Surface fish used in this experiment are derived from a lineage that is more closely related to the Molino cave fish population than to Tinaja and Pachón (23). Brains were dissected from adult animals, serial sectioned at 8 µm thickness, stained with cresyl violet dye (Nissl), and imaged, resulting in 424-760 sections per brain. We then registered all brain slices such that they aligned with one another, and imported the data into AMIRA 3D rendering software, where serial-sections were volumetrically reconstructed to generate a 3-dimensional brain (Figure 1B, Supplemental Movie 1-4).

**Figure 1.**
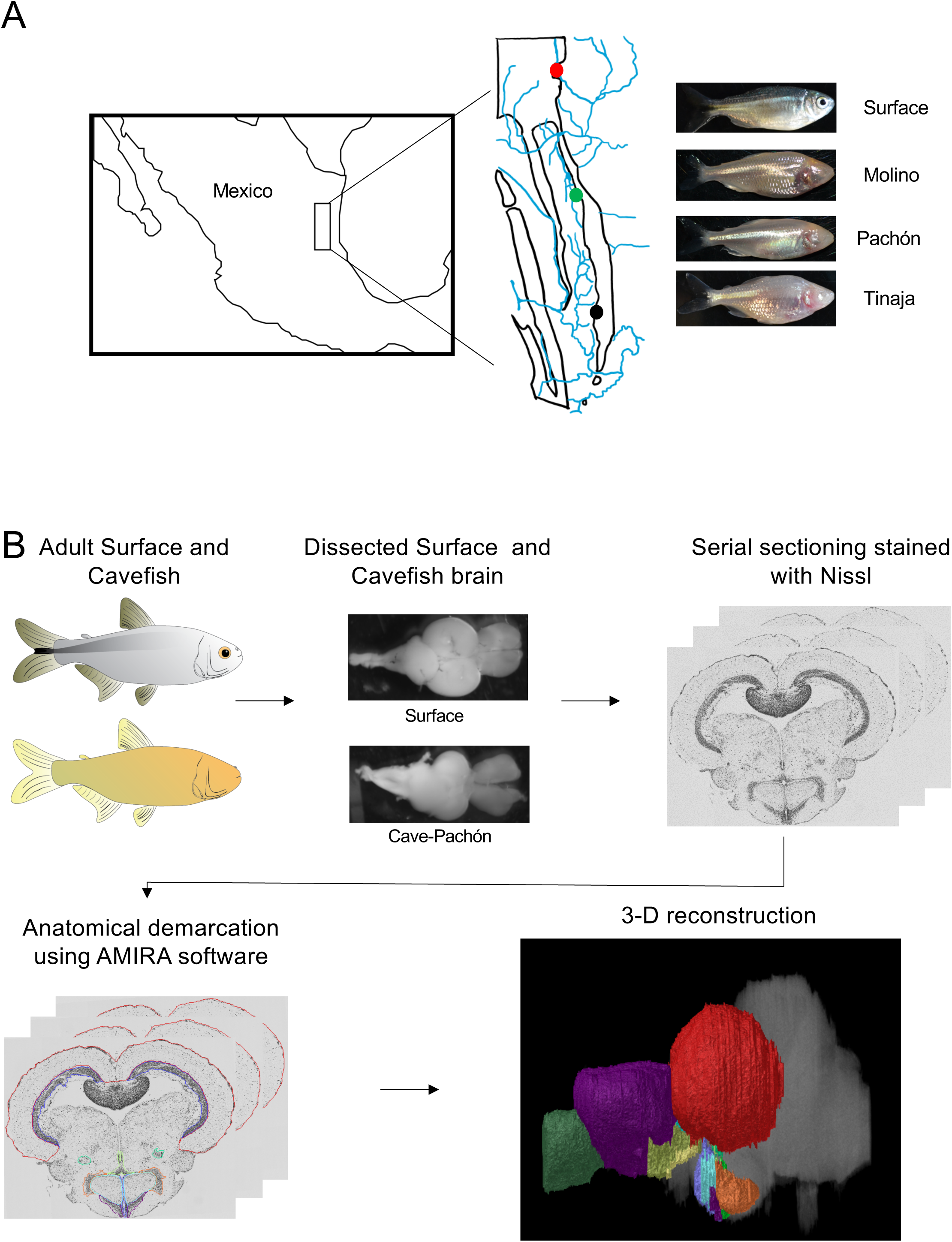
Overview of experimental design. (A) Map of Mexico with location of Molino (red), Pachón (green) and Tinaja (black) caves. (B) Flow chart diagraming experimental design and procedure.

Specific neuroanatomical regions in each brain were identified by comparing an adult zebrafish brain atlas (24), and a previously annotated brain of the cavefish from the Micos cave (25), a hybrid cave population (population with cave and surface-like animals) of the new lineage (22). After locating individual neuroanatomical regions, we defined each brain nucleus by demarcating the boundaries of the region throughout serial sections using AMIRA. We then quantified a volume of each region (Figure 1B). The volume of each quantified region was normalized to the total brain volume, measured from the anterior telencephalon to the posterior cerebellum providing a measurement of relative volumetric enlargement or reduction in size between *A. mexicanus* populations.

### Regression of optic tectum volume in cavefish populations

As proof of principle, we first quantified the optic tectum cell body (Figure 2A-C, red) and neuropil (Figure 2A-C, blue) layers, which have been reported as reduced in Pachón cavefish (20,26). The optic tectum in adult teleosts is a laminated structure with a dense region termed the optic neuropil lying ventral to the tectum. In agreement with previous findings, whole-brain reconstructions revealed a nearly two-fold reduction in tectum size in Pachón cavefish populations, as well as Molino and Tinaja populations, compared to surface fish (Figure 2A-C). To increase statistical power, we combined the total optic tectum volume of all cave populations and compared to surface fish. This comparison revealed significant differences in volume between surface and cave morphs (Figure 2D). Quantification of total volumes between surface and the three cave populations revealed a substantial reduction in total volume (Pachón = 68.69% decrease in volume compared to surface fish, Tinaja = 67.68% decrease in volume compared to surface fish, Molino = 50.51% decrease in volume compared to surface fish). In addition to the cell body layer of the tectum, the volume of the optic neuropil appeared qualitatively reduced across all three cave populations as well (Figure 2C), and quantification of volumes showed that the neuropil was smaller in cavefish relative to surface animals (Pachón = 43.24% decrease in volume compared to surface fish, Tinaja = 35.14% decrease in volume compared to surface fish, Molino = 24.32% decrease in volume compared to surface fish). When all three cavefish populations are clustered together, the optic neuropil size was reduced significantly compared to surface fish (Figure 2E). These findings extend previous observations in Pachón cavefish to Molino and Tinaja (20), revealing convergence on reduced size of the optic tectum in adult cavefish populations.

**Figure 2.**
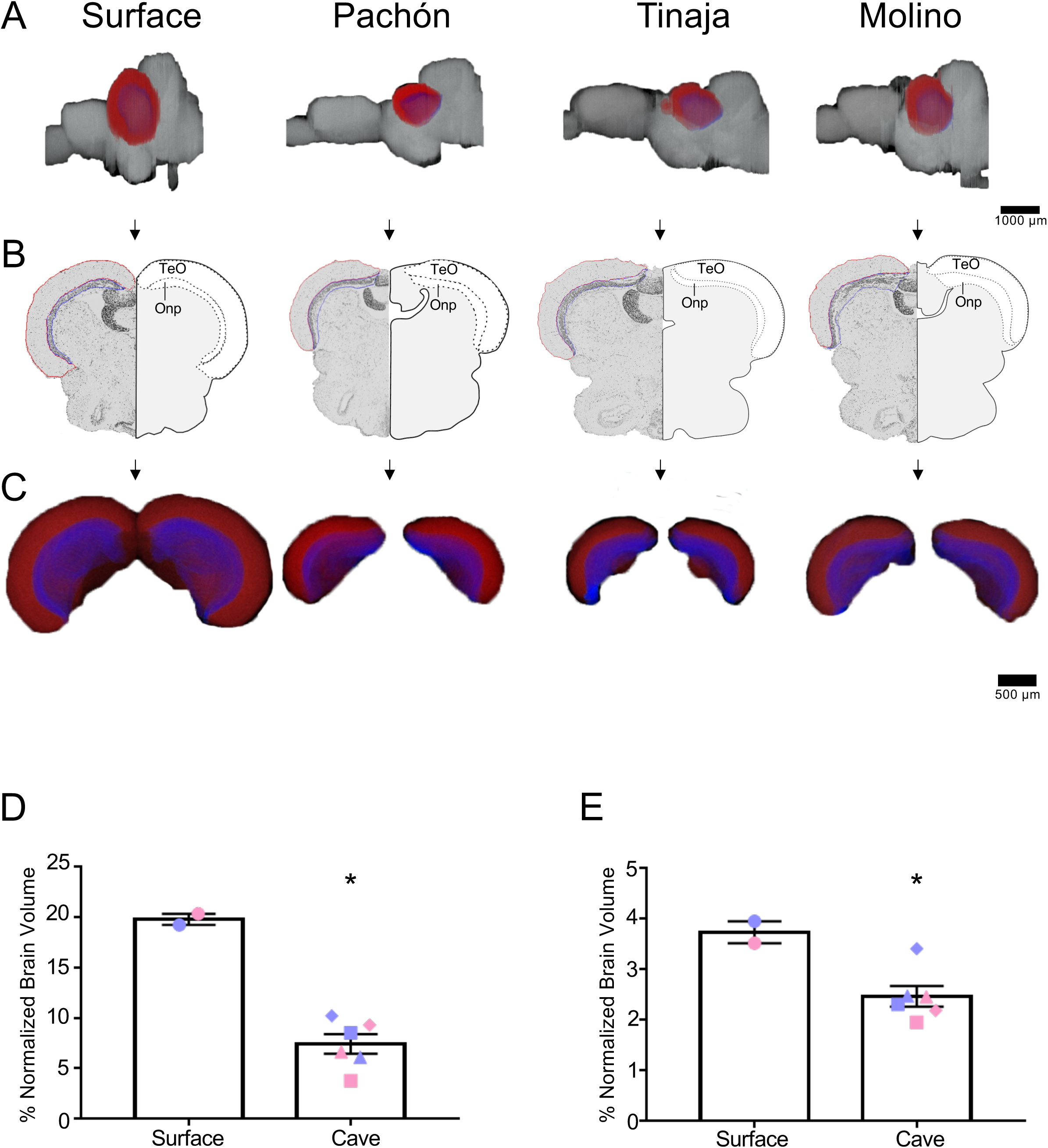
Three-dimensional reconstruction reveals regression of the optic tectum. (A) 3-D reconstructions of Surface, Pachón, Tinaja and Molino with demarcated optic tectum (red) and optic neuropil (blue) displayed. (B) Images of demarcated sections that were Nissl stained (left) and a cartoon of demarcated region (right). (C) 3-D reconstruction of the optic tectum and optic neuropil only, displayed from an anterior view. (D) Quantification of the volume of optic tectum, normalized to the size of the total brain, for surface fish and for the three cave populations. In order to examine significance, volumetric data for cavefish brains were pooled. Optic tectum of cavefish was significantly smaller than those of surface fish conspecifics (surface fish 19.8 ± 0.55; cavefish 7.4 ± 0.97; t-test t=6.921, p<0.05). (E) Quantification of the volume of optic neuropil. The optic neuropil was also significantly smaller in cavefish, compared to surface conspecifics (surface fish 3.7 ± 0.22; cave fish 2.5 ± 0.21; t-test t=3.272, p<0.05). Graphs in D and E are the mean ± standard error of the mean. Asterisk represent significance below p = 0.05. Blue shapes on bar graphs denote males, whereas pink denotes female. Square points on graphs represent Pachón, triangle points on graphs represent Tinaja and diamond points on graphs represent Molino.

### Expansion of the telencephalic nuclei in cavefish populations

The telencephalon modulates diverse behaviors that differ between surface and cavefish, including sleep, stress, and aggression (11,27–30). We therefore quantified telencephalon volume across *A. mexicanus* populations and found it to be expanded in all three populations of cavefish compared to surface fish (Figure 3A-C). Comparing total volume for surface fish and the combined data for cavefish populations revealed a significant increase in volume in cavefish (Figure 3D; Pachón = 52.11% increase in volume compared to surface fish, Tinaja = 57.04% increase in volume compared to surface fish, Molino = 57.75% increase in volume compared to surface fish). Despite these cave populations representing independent origins of the cave morph, no qualitative differences were observed between cavefish populations, suggesting that evolution repeatedly shapes brain morphology in similar ways. In addition, we observed differences in telencephalon shape between surface and cavefish populations. In all three cavefish populations the telencephalon is longer along the anterior-posterior axis than in surface fish (Figure 3A). Collectively, these data reveal a robust expansion of the telencephalon across three independent cavefish populations.

**Figure 3.**
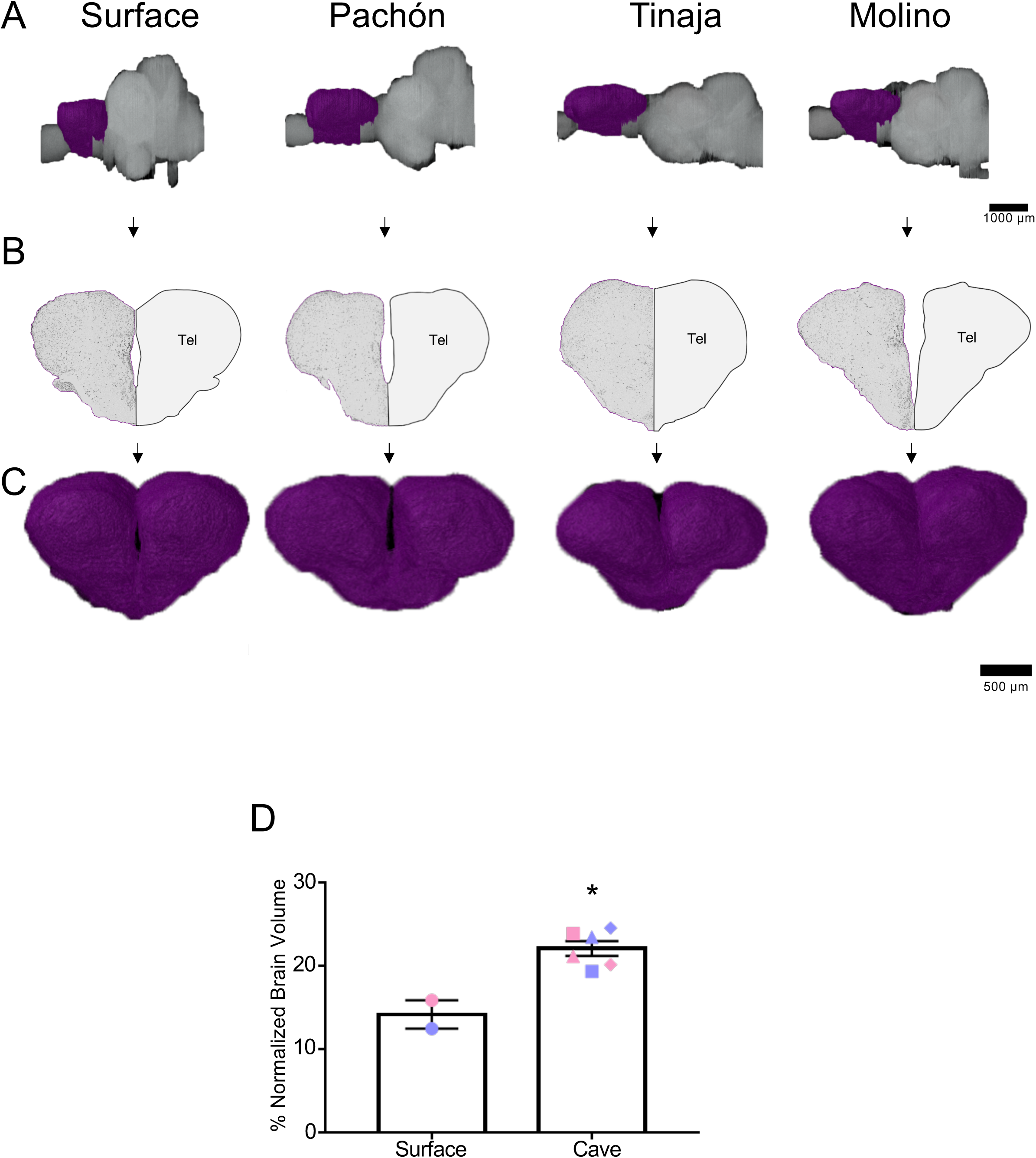
Expansion of telencephalon in cavefish populations. (A) 3-D reconstructions of Surface, Pachón, Tinaja, and Molino with demarcated telencephalon (purple) displayed. (A) Images of demarcated sections that were Nissl stained (left) and a cartoon of demarcated region (right). (C) Close up view of the 3-D reconstruction of the telencephalon from an anterior view. (D) Quantification of telencephalic volume shows expansion of the forebrain in cavefish (surface fish 14.19 ± 1.7; cavefish 22.13 ± 0.89; t-test t=4.397, p<0.05). Graph in D is the mean ± standard error of the mean. Asterisk represent significance below p=0.05. Blue points on bar graphs denote males, whereas light red denotes female. Square points on graphs represent Pachón, triangle points on graphs represent Tinaja and diamond points on graphs represent Molino.

### Analysis of thalamic and habenula nuclei

The thalamus is a central relay unit connecting the forebrain with downstream mid- and hindbrain targets, and different regions of the thalamus have been shown in mammals to modulate diverse behaviors including stress, aggression, and sleep (31–36). Moreover, anatomy and function of thalamic nuclei are conserved among mammals and fish (37–40). Quantification of the entire thalamus revealed no significant differences in gross volume cave and surface fish (Figure 4A-D; Pachón = 3.57% increase in volume compared to surface fish, Tinaja = 25.0% increase in volume compared to surface fish, Molino = 53.0% increase in volume compared to surface fish). We then examined volumetric differences between thalamic subnuclei, including the posterior, anterior, ventrolateral, ventromedial, intermediate, and central posterior (Figure S1). Of these, the posterior thalamic nuclei were significantly larger in the cavefish populations (Figure S1A). The anterior, ventrolateral and ventromedial thalamic subnuclei were, on average, larger in cavefish populations, with volumetric differences approaching significance (Figure S1B-C). By contrast, no differences were observed for the intermediate and central posterior nuclei (Figure S1D-E).

**Figure 4.**
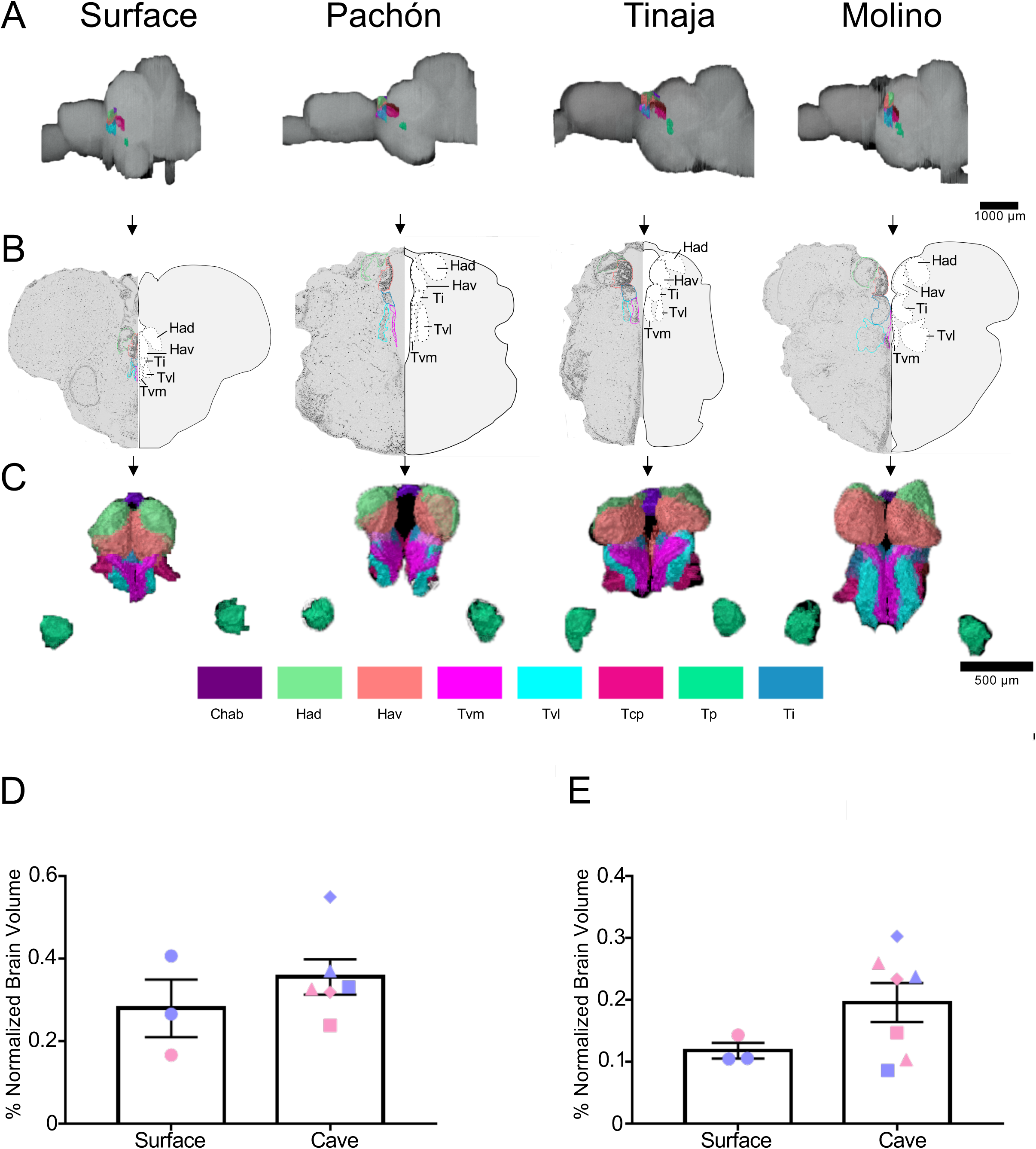
Quantification of the thalamus and habenulae reveals difference in some, but not all, subnuclei. (A) 3-D reconstructions of Surface, Pachón, Tinaja, and Molino with demarcated thalamus and habenula displayed. (B) Images of demarcated sections that were Nissl stained (left) and a cartoon of demarcated region (right). (C) 3-D reconstruction of the thalamus and habenulae only, displayed from an anterior view. (D) Quantification of the total volume of the thalamus revealed no significant differences (surface fish 0.28 ± 0.07, cavefish 0.356 ± 0.04, t-test t=0.9828, p=0.3585). (E) Quantification of the total volume of the habenulae revealed no significant differences. (surface fish 0.12 ± 0.013, cavefish 0.20 ± 0.03, t-test t=1.545, p=.1609). Graphs in D and E are the mean ± standard error of the mean. Asterisk represent significance below p=0.05. Color key for subnuclei (Chab= Habenula Commissure, Had= Dorsal Habenular Nucleus, Hav= Ventral Habenular Nucleus, Tvm= Ventromedial thalamic nucleus, Tvl= Ventrolateral thalamic nucleus, Tcp= Central posterior thalamic nucleus, Tp= Posterior thalamic nucleus, Ti= Intermediate thalamic nucleus). Blue points on bar graphs denote males, whereas light red denotes female. Square points on graphs represent Pachón, triangle points on graph represent Tinaja and diamond points on graph represent Molino.

The habenular nuclei are a conserved brain nucleus that also connect forebrain to midbrain (41,42). In rodents and other mammals, the habenulae have been shown to regulate diverse behaviors, including sleep, stress, feeding, and social interactions (43–47). Recently, the habenular nuclei have also been found to modulate similar behaviors in zebrafish (38,40,48). Because many of the behaviors modulated by the habenulae differ between surface and cave morphs, we examined volumes of the subnuclei of the habenulae. The habenula is comprised of the dorsal and ventral habenula, and its commissure (49), and this neuroanatomy is conserved among vertebrates (50). The entire habenulae was enlarged in most cavefish populations, though this did not reach statistical significance (Figure 4A-C, E; Pachón= 0.0% increase in volume compared to surface fish, Tinaja = 66.0% increase in volume compared to surface fish, Molino = 125.0% increase in volume compared to surface fish). Examining individual subnuclei revealed an expansion of dorsal habenular nucleus (Had) and ventral habenular nucleus (Hav) (Figure S2A,B) across all three cavefish populations. The habenular commissure (Chab) also appeared enlarged cave populations although there was a high-level of inter-animal variability (Figure S2C). Taken together, these findings reveal subnuclei-specific differences within the habenula of cavefish.

### Analysis of the hypothalamus reveal evolutionary changes to some but not all subnuclei

The hypothalamus controls numerous homeostatically regulated behaviors that are known to differ between surface fish and cavefish, including sleep, feeding, stress, and social behaviors (10,15–18,51). To determine whether these behavioral changes are accompanied by alterations in anatomy, we quantified the overall size of the hypothalamus, as well as individual subnuclei that modulate distinct behaviors in mammals (Figure 5 and Figure S3). We found the total volume of the hypothalamus was enlarged across all three cavefish populations compared to surface fish (Figure 4A-D, Supplemental Movie 5-8; Pachón = 41.78% increase in volume compared to surface fish, Tinaja = 36.44% increase in volume compared to surface fish, Molino = 48.0% increase in volume compared to surface fish). An expanded hypothalamus in cavefish has been demonstrated previously for larval forms (19), and thus these data reveal that hypothalamic expansion is conserved through adulthood.

**Figure 5.**
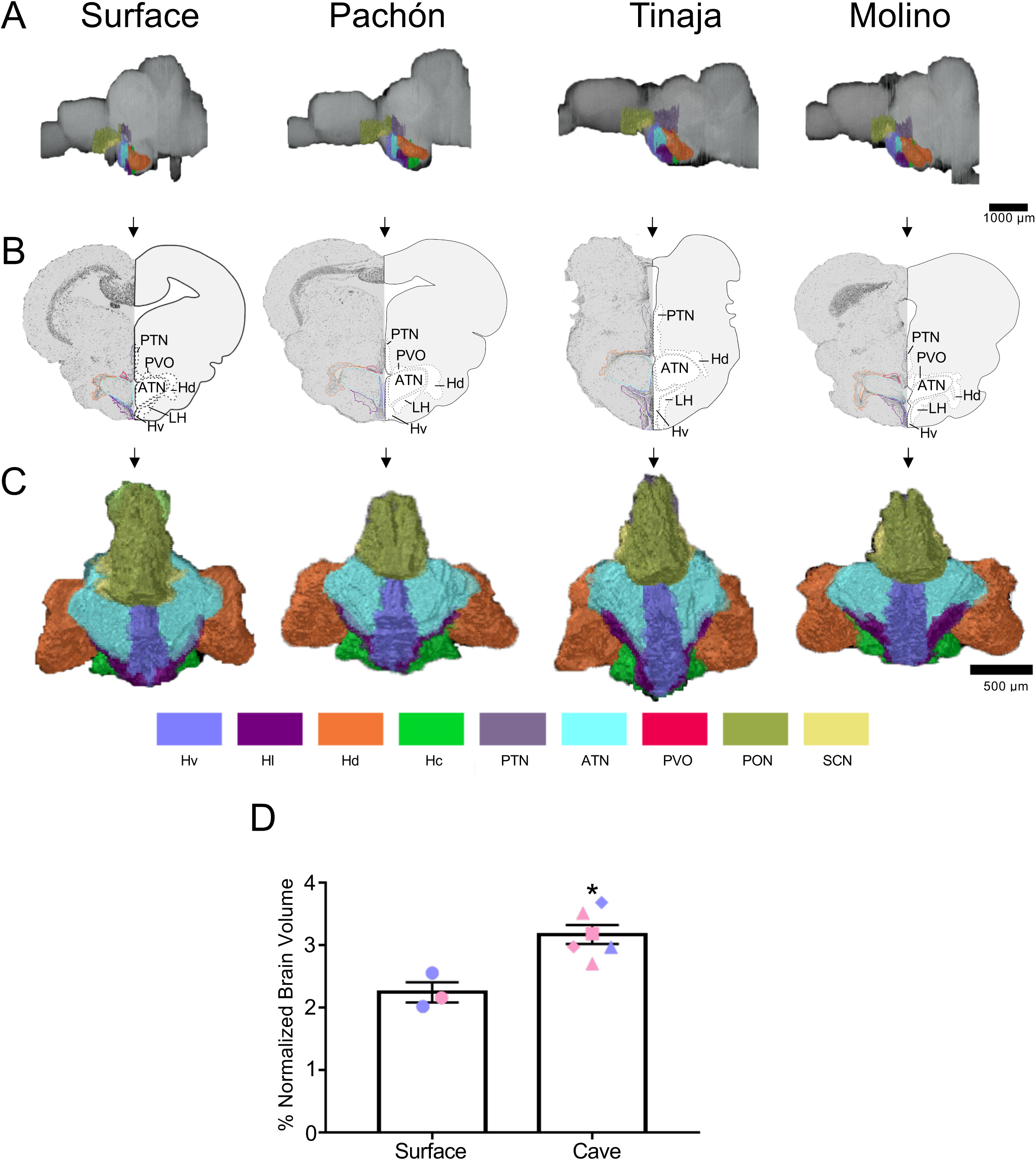
Quantification of the hypothalamus reveals expansion in cave populations. (A) 3-D reconstructions of Surface, Pachón, Tinaja and Molino with demarcated hypothalamus displayed. (B) Images of demarcated sections that were Nissl stained (left) and a cartoon of demarcated region (right). (C) 3-D reconstruction of the hypothalamus only, displayed from an anterior view. (D) Quantification of the total volume of the hypothalamus revealed significant differences (surface fish 2.25 ± 0.16, cavefish 3.18 ± 0.15, t-test t=3.802, p<0.05). Graph in D is the mean ± standard error of the mean. Asterisk represent significance below p=0.05. Color key for subnuclei (Hv= Ventral hypothalamus, Hl= Lateral hypothalamus, Hd= Dorsal Hypothalamus, Hc= Caudal hypothalamus, PTN= Posterior tuberculum, ATN= Anterior tuberculum, PON= Paraventricular organ, SCN= Superchiasmatic nucleus). Blue points on bar graphs denote males, whereas light red denotes female. Square points on graphs represent Pachón, triangle points on graphs represent Tinaja and diamond points on graphs represent Molino.

We next examined the volume of different hypothalamic subnuclei (Figure S3). We first measured the suprachiasmatic nucleus (SCN). The SCN is a critical regulator of circadian rhythms in mammals (52,53). SCN size was reduced in Pachón cavefish relative to surface fish (though interpretations are limited by the small sample size) but did not differ between surface fish and Molino or Tinaja cavefish, suggesting cave-population specific differences in SCN anatomy (Figure S3A). Conversely, the size of the following three hypothalamic subnuclei were enlarged: lateral hypothalamus (Hl), dorsal hypothalamus (Hd) and caudal hypothalamus (Hc) across cavefish populations compared to their surface conspecifics (Figure S3B-D). When all cavefish individuals are combined, these nuclei are significantly enlarged in cavefish. Furthermore, ventral hypothalamus (Hv, p = 0.077), paraventricular hypothalamus (PVO, p = 0.071), and the preoptic nucleus (PON, p = 0.074) were larger in cavefish relative to surface fish, with volumetric differences between surface and cave animals approaching significance (Figure S3E-G). These findings reveal that differences in specific subnuclei, and not overall enlargement of the hypothalamus is responsible for the observed difference in size between surface fish and cavefish.

## Discussion

Here we present an adult brain atlas for surface *A. mexicanus* and three populations of cavefish. A highly detailed brain atlas has been previously generated in zebrafish (24), and another brain atlas has been published in a cave/surface hybrid population of *A. mexicanus* cavefish (though it is untranslated from German) (25). These two resources provide a point of comparison for identifying neuroanatomical loci in cave and surface populations of *A. mexicanus*. An estimated ~100-250 million years ago of divergence separate *A. mexicanus* and *Danio rerio* (54,55). We found the brains of *A. mexicanus* were largely homologous to zebrafish, allowing for identification major brain structures.

Our analysis provides the first comparative brain atlas for surface and cave populations of *A. mexicanus*. The use of automated serial sectioning allows for volumetric reconstruction of brain regions and semi-quantitative comparisons of neuroanatomy between surface and cavefish populations. While this approach is technically feasible, practically it is limited due to the labor-intensive nature of manually tracing brain regions, and difficulties obtaining complete sectioned brains. In this study, we chose to focus on the visual system as a proof-of-principle, as well as the hypothalamus, thalamus, and habenula due to their known role in behavioral regulation. While the small number of replicates largely prevented statistical comparisons between individual cavefish populations, the robust volume differences observed between surface and cave populations for many brain regions suggest this approach may be practical for detailed anatomical comparison. Here, we have made all raw data available so that others may quantify additional brain regions of interest.

Brain atlases have been widely used in a number of species, including zebrafish, and have expanded greatly our understanding of how individual neuronal areas modulate myriad behaviors (24,56–61). Brain atlases have been generated in larval zebrafish that provide near single-neuron resolution of brain structures (56,57,62,63). The transparency of the zebrafish larvae allows for the application of functional imaging approaches (64,65), that can then be mapped on brain atlases to identify changes in activity within defined neurons (62). *A. mexicanus* larvae, like zebrafish, are transparent, providing potential for the generation of a high-resolution brain atlas.

While this level of accuracy is not possible in adult fish, due to the larger size of the brain and the need for sectioning, the added complexity of the adult brain and its considerable homology to the mammalian brain is particularly effective in comparative neuroanatomy. Further, a number of behaviors that differ between surface and cave individuals are not present in larval forms. For example, a loss of aggressive behavior has been documented in cavefish animals (11), and other studies have demonstrated that cavefish do not school, whereas their surface conspecifics do (16). Many of these behaviors are not present in larval forms, and thus an adult atlas facilitates identification of brain regions that modulate more complex behaviors only seen in adults.

In this study, brain regions were standardized to the overall size of the brain from the anterior telencephalon to the posterior cerebellum since these areas were the most consistent between samples. To correct for individual differences in size and growth rate, we normalized all brain volumes (66). Quantitative comparisons between smaller neuroanatomical regions, such as subnuclei within the hypothalamus or thalamus, may be confounded by large differences within other brain regions, such as the optic tectum. However, the variability in differences between subnuclei suggests localized changes in brain volume can be detected. As an example, most nuclei in the hypothalamus are expanded across cavefish populations, yet no differences are detected within the SCN for Tinaja and Molino caves relative to surface.

Our findings identify the expansion of multiple hypothalamic nuclei, suggesting shared processes may govern evolved differences in hypothalamic development. The hypothalamus in cavefish larvae is expanded through a mechanism that is dependent on the differential expression of several morphogens and transcription factors, including sonic hedgehog and Nkx2.1 (19). One hypothesis is that reduced anatomical constraints from eye-loss allow for hypothalamic expansion. A number of hypothalamic neuropeptides are known to be upregulated in cavefish including HCRT and NPY, which localize to the lateral hypothalamus and periventricular/lateral hypothalamus respectively (7,8,67). Both of these nuclei are larger across all three populations of cavefish. Many hypothalamus-regulated behaviors including sleep, feeding, aggression, and sociality are altered in cavefish (8,10,11,15–17,68), suggesting hypothalamic function may be a under significant selective pressure.

In agreement with the previous literature, we identify convergent evolution of changes in brain regions associated with sensory processing (20,26,69). The optic tectum is significantly reduced across all three cavefish populations. These findings are consistent with an increased reliance on non-visual cues in cave animals (9,70). Future work will explore how the reliance on nonvisual cues has shaped brain regions. For example, taste buds are more numerous in cavefish (69,71) and the lateral line of cavefish is also significantly expanded, suggesting increased reliance on sensory processes that do not involve sight (72,73). The sensory neurons from taste and mechanosensation neurons project to the nucleus of the solitary tract (NST) and medial octavolateralis nucleus (MON) within the brain, respectively (74–76). Based on findings from other sensory pathways, these regions may be predicted to be enlarged. Future analysis of serially sectioned brains will allow for detailed quantification and comparison of sensory structures between *A. mexicanus* populations.

Here, we used brains stained with Nissl, and demarcated manually individual regions of the adult brain. We see two main future expansions of this work. First, future efforts will streamline the labor-intensive approach of manual demarcation of individual regions. Similar large-scale neuroanatomical reconstruction efforts, such as electron microscopy tracing of the *Drosophila* brain have been successful in analyzing large data sets like these (77). It is also possible that automated tracing methodology may be developed to reduce the time required for analysis. Further, future imaging of additional serially-section brains may allow for more quantitative comparisons between populations. Second, in zebrafish and other models, transgenic labeling of precise neuronal population has facilitated greatly the demarcation of individual neuronal regions (78,79). Moreover, transgenic labeling of neurons in the brain permits tracing of neuronal projection, something that is not possible with Nissl staining (68,80). Whereas transgenic technology has not been widely used in *A. mexicanus*, recent studies have shown that the Tol2 system, which is widely used in zebrafish, is highly effective in *A. mexicanus* surface and cavefish (81–83). Future work incorporating these tools would facilitate a highly defined neuroanatomical brain atlas for the *A. mexicanus* adult brain.

## Methods

### Fish husbandry

Animals care and husbandry were carried out as previously described (8,84). Briefly, adult *A. mexicanus* stocks were originally obtained from the Jeffery (University of Maryland) or Borowsky (New York University). These fish have been bred and maintained on a recirculating aquatics system at Florida Atlantic University. The water temperature was maintained at 21 ± 1°C, and the lights were maintained on a 14:10 LD cycle (25-40 lux at lights on). All fish were fed a mix of fish flakes (TetraMin) and California black worms (Aquatic Foods). All experiments in this study were approved by the Institutional Animal Care and Usage Committee (IACUC) at Florida Atlantic University, protocol numbers A17–21 and A15–32, or the IACUC at Stowers Institute for Medical Research. All fish used in this study were approximately 1 year old. A total of 10 brains were dissected and analyzed per population. We used 2 male and 1 female brains from surface population, 1 male and 1 female brains from Pachón population, 1 male and 2 female brains from Tinaja population, and 1 male and 1 female brains from Molino population. In some cases, brains could not be quantified for all neuroanatomical regions due to tissue damage.

### Sectioning

Fish were euthanized by incubation in MS-222 (500 mg/mL) for 10 min and decapitated using sharp scissors. The head was immediately fixed with freshly prepared 4 % paraformaldehyde (PFA, diluted from 16% (wt/vol) aqueous solution, Electron Microscopy Sciences, cat# 15710) in 1 x PBS for 48 hours at 4 ° C with a change of 4 % PFA / 1xPBS after 3 hours. Heads were washed three times in 1xPBS and subsequently, brains were dissected according to Moran et al. (26). Brains were dehydrated through graded ethanol (30%, 50%, 70%) and processed with a PATHOS Delta hybrid tissue processor (Milestone Medical Technologies, Inc, MI) followed by paraffin embedding. Coronal slices of paraffin sections with 8 µm thickness were continuously cut using a Leica RM2255 microtome (Leica Biosystems Inc. Buffalo Grove, IL) and mounted on Superfrost Plus microscope slides (cat# 12-550-15, Thermo Fisher Scientific). Nissl staining was performed as described in Vacca et al. (85). Briefly, sections were deparaffinized and hydrated to distilled water. Sections were stained in cresyl echt violet (0.5 g cresyl echt violet (CI 51010); 80 mL distilled water; 20 mL absolute alcohol) for 8 minutes, briefly rinsed in distilled water, dehydrated with 95 % absolute alcohol 2 times, subsequently cleared in 2 changes of xylene and finally mounted. Slides were scanned using an Olympus slide scanner VS120 with a 20x objective. Images were extracted from VSI files in sequence using a customized plugin in Fiji (ver 1.51H)(86), a mask constructed, and registered using a multithreaded version of StackReg1 (87). Blank spaces in the registered image were filled with artificial noise that matched the all-white background using a custom plugin in Fiji. Plugins are available at https://github.com/jouyun/smc-plugins and https://github.com/cwood1967/IJPlugins/

### Volumetric Reconstruction

ImageJ FIJI (ver 1.51H)(86) was used to convert serial sections to a .tif image sequence. Image sequence was uploaded into the AMIRA software (ver 6.2.0, Thermo Fisher, Waltham, MA). To create proper demarcations, neuroanatomical regions of interest (ROIs) from Nissl stains were set under the “segmentation” tab using the lasso tool. To view 3-dimensional reconstructions of neuroanatomical ROI’s, a ‘volren’ object was created under the “project” tab. Volren object was connected to the original .tif image sequence as well as the label fields used to create demarcated neuroanatomical ROI’s.

### Measurements and Statistical analysis

To quantify total volume of induvial demarcated regions (i.e., each ROI), we used the ‘volume per VOI’ result of the ‘material statistics’ function in AMIRA (ver 6.2.0). To correct for differences in size and growth rate among different fish populations, all volumetric results were normalized to the length of the brain from the anterior telencephalon to the posterior cerebellum; volumetric measurements were thus calculated as a percentage of volume relative to this normalized length. For statistical comparisons of ROI volumes between two groups (i.e. the pooled cavefish data compared to surface), we used a standard two-tailed t-test. For statistical comparisons of more than 2-groups, a parametric ANOVA was implemented. All statistics were performed using GraphPad Prism (ver 7.0).

## Supporting information

Supplemental Movie 1, Whole Brain of surface

Supplemental Movie 2, Whole Brain of Pach??n

Supplemental Movie 3, Whole Brain of Tinaja

Supplemental Movie 4, Whole Brain of Molino

Supplemental Movie 5, Hypothalamus of surface

Supplemental Movie 6, Hypothalamus of Pach??n

Supplemental Movie 7, Hypothalamus of Tinaja

Supplemental Movie 8, Hypothalamus of Molino

## Acknowledgements

The authors would like to acknowledge Nancy Thomas and the histology core at the Stowers Institute for support on the brain sectioning. Furthermore, we would like to thank the aquatics team at the Stowers Institute for husbandry of the fish. This work was supported by grants R15MH118625 to E.R.D., R21NS105071 to E.R.D and A.C.K., R01GM127872, NSF IOS165674, BSF2018-190 to A.C.K, and by institutional funding to NR. NR was supported by the Edward Mallinckrodt Foundation and JDRF. RP was supported by a grant from the Deutsche Forschungsgemeinschaft (PE 2807/1-1).

## Data availability statement

Original data underlying this manuscript can be accessed from the Stowers Original Data Repository at http://www.stowers.org/research/publications/libpb-1427. All original and analyzed data will also be provided upon request.

---

**Supplemental Figure 1.**
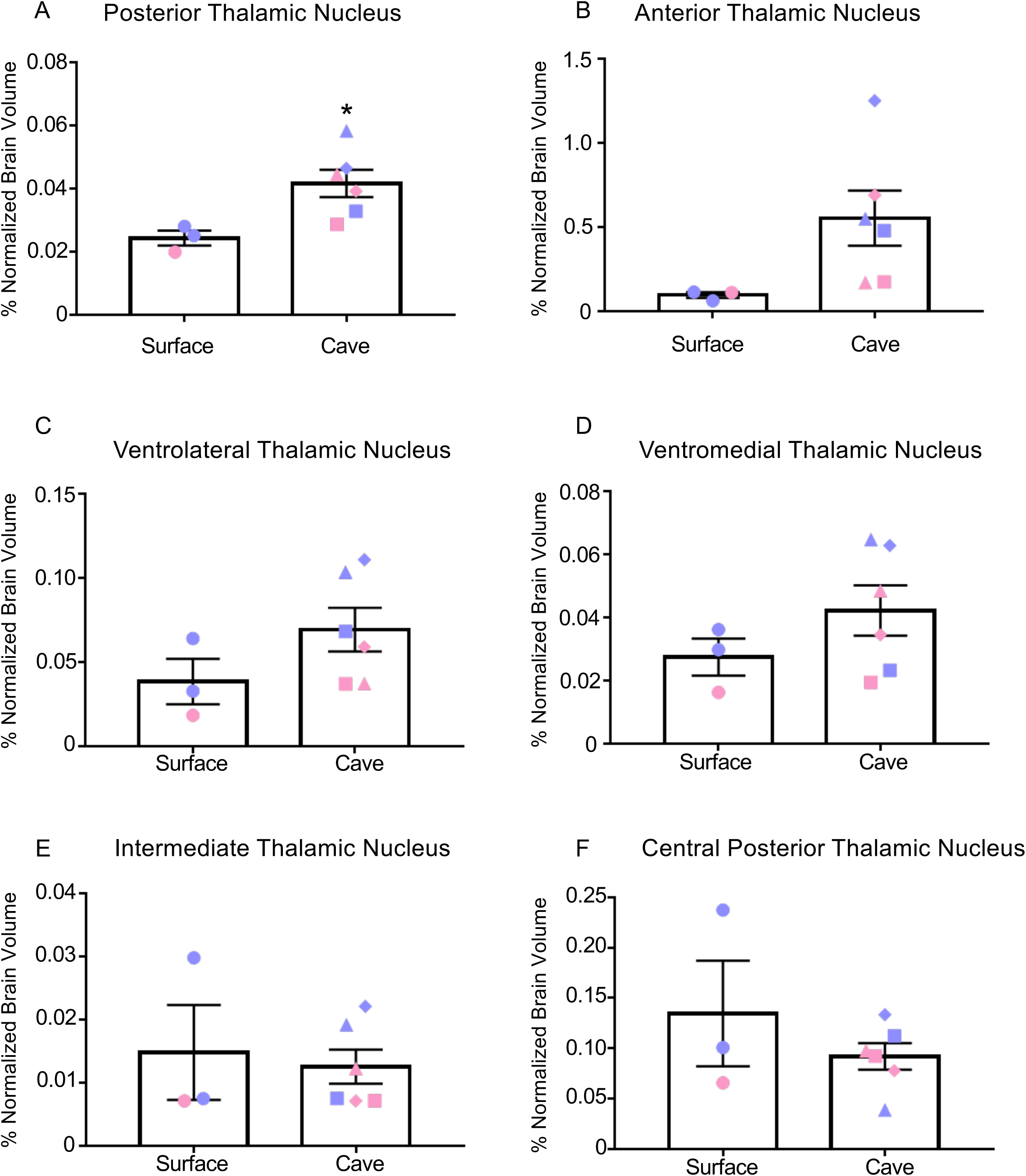
Analysis of different thalamic subnuclei reveals expansion of some, but not all, regions. (A) Quantification of the posterior thalamic nucleus shows an expansion of volume (surface fish 0.02 ± 0.023, cavefish 0.042 ± 0.004, t-test t=5.656, p = 0.032). (B-C) analysis of volumes from anterior thalamic nuclei (B) and ventrolateral thalamic nucleus (C) revealed difference that approached significance (Ta = surface fish 0.010 ± 0.001, cavefish 0.06 ± 0.02; t-test t=1.907; p=0.098; Tvl = surface fish 0.04 ± 0.01, cavefish 0.07 ± 0.013; t-test t=1.474; p=0.184). (D-F) There were no significant difference for ventromedial thalamic nucleus (D), intermediate thalamic nucleus (E), and central posterior thalamic nucleus (F) (Tvm =surface fish 0.03 ± 0.006; cavefish 0.04 ± 0.008; t-test t=1.203; p=0.268; Ti = surface fish 0.015 ± 0.008; cavefish 0.013 ± 0.003; t-test t=0.3565, p=0.732; Tcp = surface fish 0.134 ± 0.05; cavefish 0.092 ± 0.013; t-test t=1.083, p=0.315) All graphs are the mean ± standard error of the mean. Blue points on bar graphs denote males, whereas light red denotes female. Square points on graphs represent Pachón, triangle points on graphs represent Tinaja and diamond points on graphs represent Molino.

**Supplemental Figure 2.**
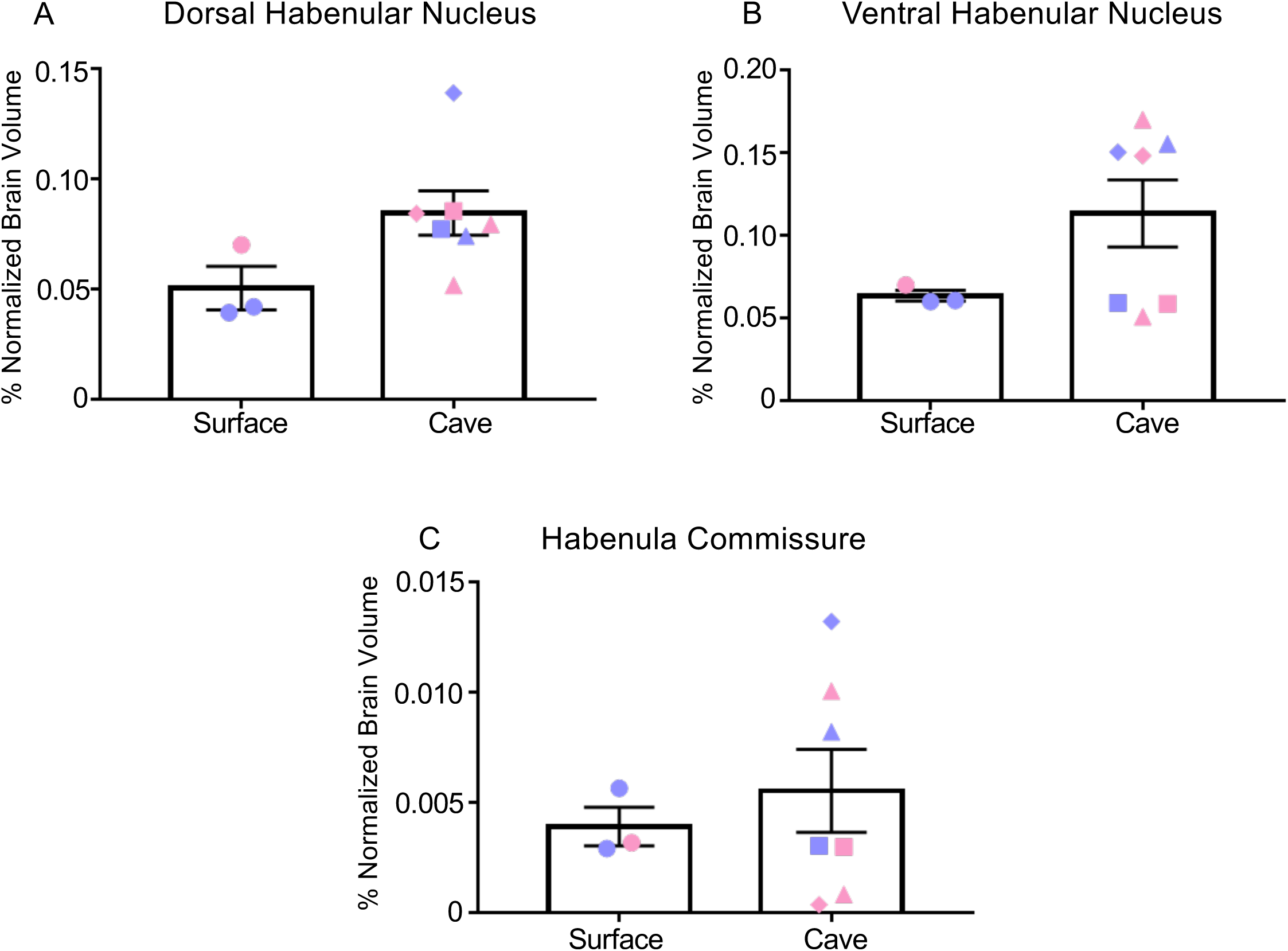
Analysis of different habenulae subnuclei reveals expansion of subnuclei, some approaching significance. (A) Quantification of the dorsal habenular nucleus shows an expansion in cavefish approaching significance (surface fish 0.05 ± 0.01, cavefish 0.08 ± 0.01; t-test t=2.01; p=0.079). (B) Quantification of ventral habenular nucleus also showed an expansion in cavefish although not statistically significant (surface fish 0.06 ± 0.003, cavefish 0.11 ±.02; t-test t=1.542; p=0.16). (C) Analysis of the habenula commissure showed no significance between morphs (surface fish 0.004 ± 0.0009, cavefish 0.057 ± 0.0019; t-test t=0.5352; p=0.61). All graphs are the mean ± standard error of the mean. Blue points on bar graphs denote males, whereas light red denotes female. Square points on graphs represent Pachón, triangle points on graphs represent Tinaja and diamond points on graphs represent Molino.

**Supplemental Figure 3.**
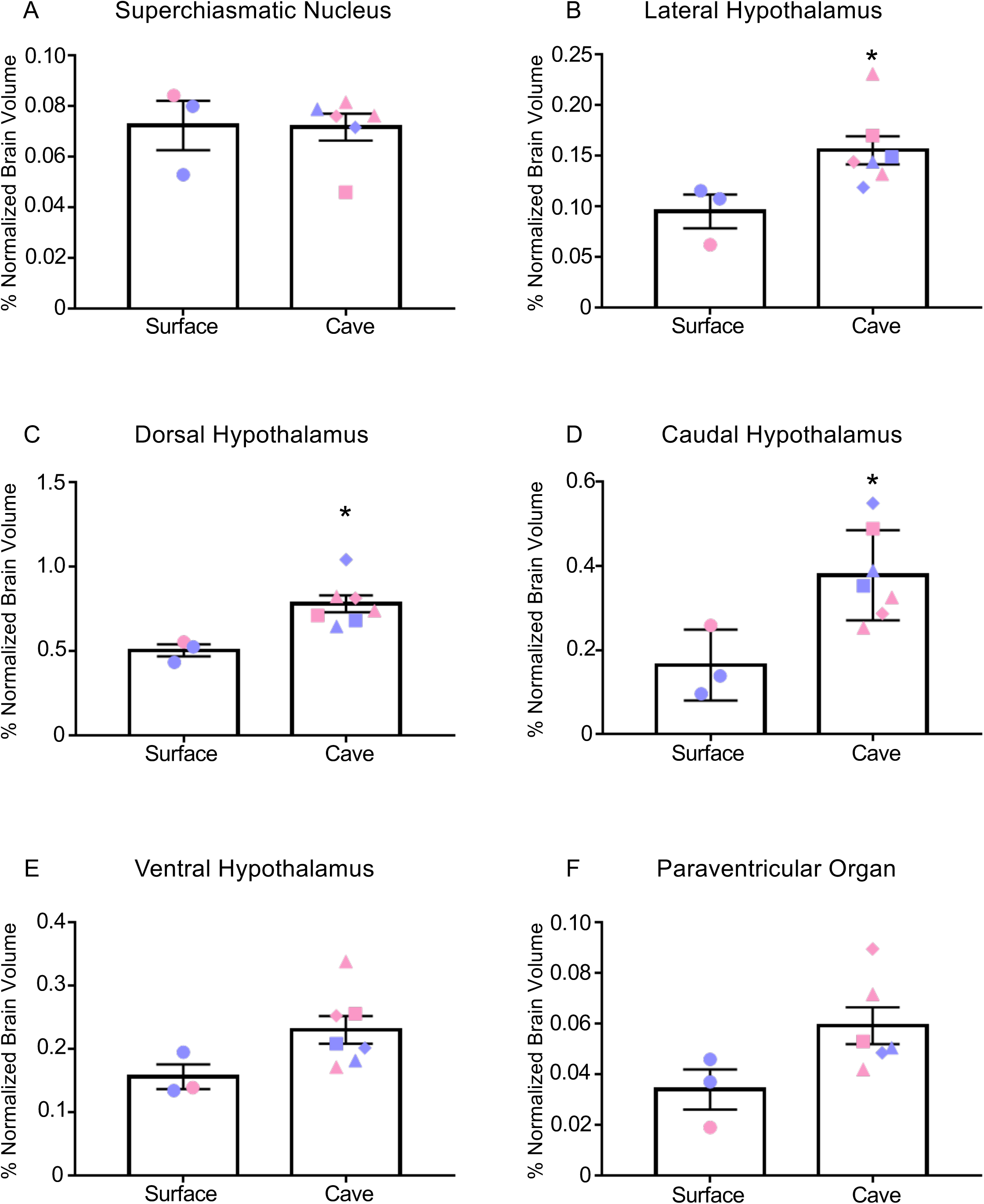
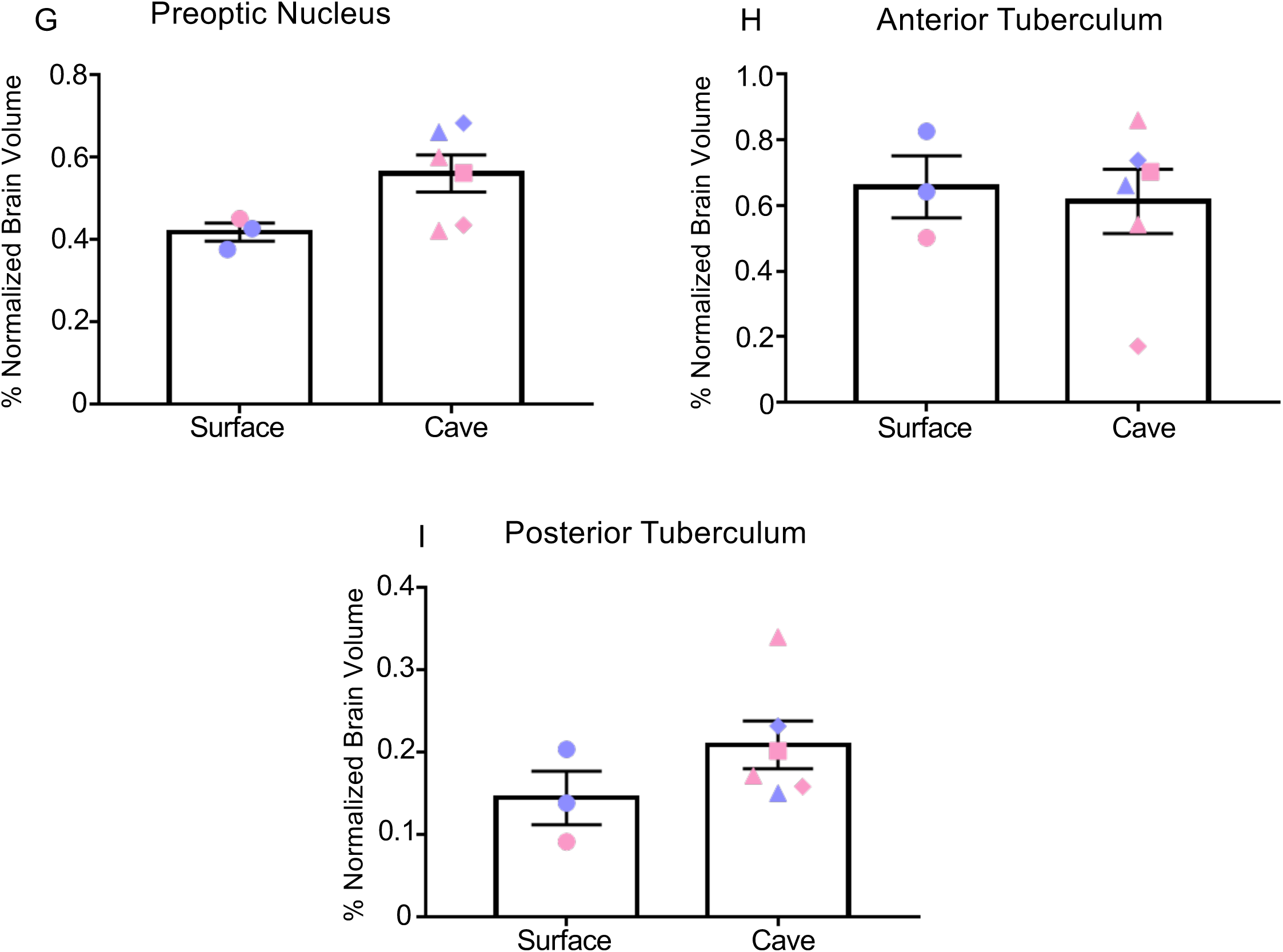
Analysis of different hypothalamic subnuclei reveals significant expansion of lateral, dorsal, and caudal hypothalamus in cavefish while others remain similar to surface fish. (A) Analysis of the superchiasmatic nucleus showed no difference between morphs (surface fish 0.07 ± 0.01, cavefish 0.072 ± 0.005; t-test t=0.0637; p=0.95). (B-D) Significant differences were observed between the lateral hypothalamus (Hl) (B), dorsal hypothalamus (Hd) (C) and caudal hypothalamus (Hc) (D) (Hl = surface fish 0.095 ± 0.04, cavefish 0.12 ±0.014; t-test t=2.506; p<0.05, Hd =surface fish 0.5 ± 0.04, cavefish 0.78 ± 0.05; t-test t=3.034; p<0.05, Hc =surface fish 0.16 ± 0.05, cavefish 0.38 ± 0.04; t-test t=3.034; p<0.05). (E-G) Analysis of the ventral hypothalamus (Hv) (E), paraventricular organ (PVO) (D) and preoptic nucleus (PON) showed an enlargement in cavefish that approached significance (Hv =surface fish 0.16 ± 0.02, cavefish 0.23 ± 0.02; t-test; p=0.07, PVO =surface fish 0.034 ± 0.008, cavefish 0.06 ± 0.007; t-test t=2.12; p=0.07, PON =surface fish 0.42 ± 0.022, cavefish 0.56 ± 0.05; t-test t=2.098; p=0.07). (H-I) Quantification of the anterior tuberculum (ATN) (H) and posterior tuberculum (PTN) (I) showed no difference between morphs (ATN =surface fish 0.66 ± 0.1, cavefish 0.61 ± 0.1; t-test t=0.2833; p=0.78, PTN=surface fish 0.14 ± 0.03, cavefish 0.21 ± 0.03; t-test t=1.357; p=0.22). All graphs are the mean ± standard error of the mean. Blue points on bar graphs denote males, whereas light red denotes female. Square points on graphs represent Pachón, triangle points on graphs represent Tinaja and diamond points on graphs represent Molino.

**Supplemental Movie 1. Three-dimensional reconstruction of whole brain from surface fish.**

**Supplemental Movie 2. Three-dimensional reconstruction of whole brain from Pachón cavefish.**

**Supplemental Movie 3. Three-dimensional reconstruction of whole brain from Tinaja cavefish.**

**Supplemental Movie 4. Three-dimensional reconstruction of whole brain from Molino cavefish.**

**Supplemental Movie 5. Three-dimensional reconstruction of hypothalamus from surface cavefish.**

**Supplemental Movie 6. Three-dimensional reconstruction of hypothalamus from Pachón cavefish.**

**Supplemental Movie 7. Three-dimensional reconstruction of hypothalamus from Tinaja cavefish.**

**Supplemental Movie 8. Three-dimensional reconstruction of hypothalamus from Molino cavefish.**

